# SigSpace: an LLM-based agent for drug response signature interpretation

**DOI:** 10.64898/2025.12.02.691945

**Authors:** Rohit Khurana, Ishita Mangla, Giovanni Palla, Siddhant Sanghi, Daniele Merico

## Abstract

Agent systems powered by large language models (LLMs) are increasingly applied in computational biology to automate analysis, integrate data, and accelerate discovery. Here, we investigate the capacity of LLM-driven agents to interpret transcriptional response signatures of drug perturbations in cancer cell lines, a task central to understanding drug mechanisms of action (MoAs) and supporting cancer drug discovery. Leveraging the Tahoe-100M dataset of 100 million transcriptomic profiles across 1,100 small-molecule perturbations and 50 cancer cell lines, we developed an LLM-based agent system, SigSpace, that processes differential gene expression signatures and generates concise, human-readable summaries of drug responses. We then tested whether blinded response signature summaries could be correctly matched to their corresponding drug identity or MoA. Our results show that LLM-generated summaries consistently outperform random baselines and that the choice of LLM model and signature score significantly influences performance. These findings highlight the potential of LLMs to enhance interpretation of complex transcriptional data to enable drug discovery. Future directions for improvement include exploring alternative response signature formats, improving summarization fidelity, benchmarking performance across different summary formats and lengths, and broadening applications to additional datasets and predictive tasks in drug discovery.

## 1 Introduction

Agent systems powered by large language models (LLMs) are emerging as transformative tools in computational biology, offering novel capabilities for database query, data integration, analysis automation, and scientific discovery acceleration. These systems combine the reasoning and adaptive decision-making of agent-based approaches with the flexible natural language understanding and generative power of LLMs, enabling dynamic interactions with complex biological datasets and workflows. By orchestrating tasks such as literature mining, experimental design, hypothesis generation, and multi-modal data interpretation, LLM-driven agents hold the potential to augment researchers’ capacity to navigate the vast and heterogeneous landscape of biological information [1, 2, 3, 4].

In this work, we evaluate the capacity of state-of-the-art LLMs to interpret the transcriptional response signatures of drug perturbations in cancer cell lines, a key task in cancer drug discovery [5]. Drug transcriptional response can inform the drug’s mechanism of action (MoA), either by similarity to well-characterized drugs or gene perturbations, or by revealing what biological pathways are perturbed. In addition, reversal of disease-associated transcriptional signatures can be used to repurpose existing drugs for new indications, illustrating one of many ways in which these responses, often multifaceted and affecting diverse genes and pathways, can be leveraged. [6]. A common framework to analyze and interpret this type of data is to first compute differential gene expression scores and then quantify enrichment of pre-defined gene sets that capture biological pathways and processes [7]. Software like Cytoscape can be used to aid result visualization and interpretation [8], but this can be unwieldy and time-consuming for high-dimensional datasets. LLMs can support this task by summarizing results in a concise and human-readable format.

Recent work has shown the tremendous promise of building agents for scientific discovery [9, 10, 11]. In the context of drug discovery, several agent systems have been proposed that aim to enhance drug discovery through automated workflows [12, 3, 13]. LLMs have shown promising performance in summarizing gene-set enrichment analysis (GSEA) results [14], generating or verifying functional gene-set membership [15], and predicting single cell genomics perturbation response for held-out genes [16]. However, to date, the ability of LLMs to parse perturbation transcriptional signatures and support drug discovery tasks such as MoA prediction has not been demonstrated.

In this work, we developed an LLM-based agent system capable of processing and reasoning across such response signatures. We leveraged the recently released Tahoe-100M dataset, which comprises 100 million transcriptomic profiles capturing the effects of 1,100 small-molecule perturbations across 50 diverse cancer cell lines [17]. Using this dataset, our system interprets perturbation response signatures and generates summaries that include potential drug MoAs. To evaluate if the generated summary correctly captures the most important features, we assessed whether the system can accurately reverse-associate blinded drug response summaries with the corresponding drug or its MoA. In particular, we evaluated the performance of two different signature scores, derived either from GSEA [18] or Vision [19], and three state-of-the-art LLMs towards these predictive tasks. Our results demonstrate that while all modeling combinations tend to outperform a random baseline in ranking both MoAs and drug identity, the choice of model and signature representation significantly impacts predictive performance. Overall, our work provides a quantitative assessment of LLMs’ ability to interpret transcriptional gene signatures in the context of target discovery and drug prioritization for cancer drug discovery.

## 2 Methods

### 2.1 Agent system overview

SigSpace ^2^ is an agent-based system used to interrogate drug response signatures derived from the Tahoe-100M dataset. It is composed of three main processing modules that operate sequentially (Figure 1, Supplementary Figure 1):

**Figure 1:**
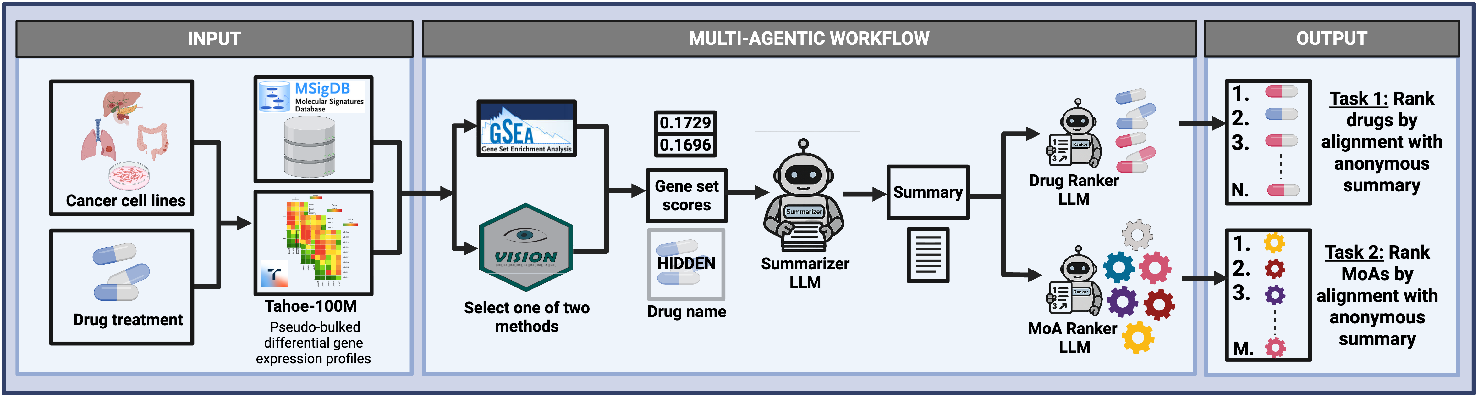
SigSpace’s agentic workflow. The system reasons over signature scores based on MSigDB gene sets, uses an LLM to generate a human-readable summary of the perturbed biological pathways, and then performs one of two tasks: drug ranking (*N* = 375) or MoA ranking (*M* = 25), where the identity of the drug used to generate the summary is hidden.

1. **Prompting:**The user specifies the drug, model, and task, and may also provide optional hyperparameters, such as particular cell lines whose metadata the system can access. This initial query defines the scope and parameters of the analysis performed by SigSpace.
2. **Signature retrieval and summarization:**The agent’s first LLM invocation is paired with tool calls that fetch precomputed perturbation response signatures from external analysis modules. These modules, implemented using Vision or GSEA, return sets of biological pathways from MSigDB [18] with associated scores. The LLM then composes a human-readable summary of the retrieved signatures to ground subsequent reasoning.
3. **Critical assessment and ranking:**The second LLM invocation performs critical assessment of the signature summary and executes one of two ranking tasks:
  a. **Drug ranking:**Using the signature summary alone, the same LLM model considers the experimental drug library from Tahoe-100M (375 compounds) and generates a ranked list of the top 50 most probable candidates, ordered from most to least likely. Drugs not appearing in the top 50 are considered to have a low probability of association with the generated summary and are thus omitted from further analysis.
  b. **MoA ranking:**Using the signature summary alone, the same LLM model analyzes defined MoAs from Tahoe-100M (25 MoAs) and ranks them from most to least likely. Drugs with an “unclear” MoA are excluded from this task, as this class includes both drugs with a well-characterized but infrequent MoA and a conflicting or uncharacterized MoA.

### 2.2 LLMs

Three LLMs were evaluated for drug ranking and MoA ranking tasks. Specifically, we used DeepSeek-R1-671B (referred to as DeepSeek-R1), Llama-4-Maverick-17B-128E-Instruct-FP8 (referred to as Llama-4-Maverick-17B), and OpenAI o4-mini (referred to as o4-mini). For each model, we generated drug rankings and MoA rankings using both GSEA and Vision scores.

### 2.3 Prompting

Although the agent can be used conversationally to explore gene signatures from the Tahoe-100M dataset, in this work, we focus on a controlled setting: evaluating how LLMs reason about perturbation transcriptional signatures. To this end, we designed a structured prompting pipeline in which the initial user query (drug, model, and task) triggers the system prompt and tool calls for signature retrieval. The drug name is used only at this stage to select the appropriate signatures, after which it is deliberately removed from the model’s context so that subsequent modules, invoking the summarization and ranking prompts, operate without direct knowledge of drug identity. This ensures that both summarization and ranking rely solely on the biological content of the signatures rather than memorized drug properties.

We also experimented with different prompt structures for the drug ranking and MoA ranking tasks. Empirically, adding domain context (e.g., describing how gene signatures were computed and how to interpret their scores) improved summary coherence and downstream ranking performance. These findings suggest that task-specific methodological detail helps constrain the model toward more accurate biological inferences. A systematic evaluation of these prompt-engineering strategies, including variations in context size and signature representation, is left for future work.

### 2.4 Perturbation response signature computation

Two sets of gene signature scores were computed from Tahoe-100M, each using MSigDB as the gene-set source:

- *Vision*: Differential Vision scores were computed by comparing the pseudobulked gene expression of drug-treated cells to plate-matched DMSO controls, further described in [17]. For LLM ingestion, we included the top 250 and bottom 250 gene sets by highest median score, after computing the median score across all cell lines at the highest drug concentration.
- *GSEA*: GSEA signatures were computed using the `fgsea` package [20], with the pseudobulked differential gene expression statistic as input, further described in [17]. We included all gene sets with an adjusted *p*-value *<* 0.05 and a minimum fraction threshold of 0.25 across cell lines, considering both positive and negative normalized enrichment scores.

When the LLM queries transcriptional signatures through tool calls, it receives a filtered list of gene sets with their corresponding scores, which form the basis for downstream summarization and ranking. For queries involving cell-line metadata, we use information from [17]. Specifically, we include the top 5 most frequently occurring cell lines from which the top and bottom scores were filtered for each gene signature tool. This approach enables us to include information only for cell lines that have reported the most significant gene signature changes under a given perturbation, thereby providing useful yet concise context for the summarization step.

### 2.5 Agent and tool design

We implemented the agent system using the LangChain framework [21], which provides utilities for integrating LLMs with external tools and APIs. Within this framework, we developed three categories of tools, each available for signatures derived from either Vision or GSEA:

- **Signature retrieval**: These tools return a set of gene signatures, filtered according to the criteria described in 2.4.
- **Signature ranking**: These tools take the LLM-generated summary of the retrieved signatures as input and produce a ranked list of either drug candidates or MoAs.
- **Cell-line metadata**: Based on the retrieved signatures, this tool injects cell-line metadata corresponding to the top 5 most frequently occurring cell lines in both the top and bottom signatures returned by the retrieval tools.

### 2.6 Model evaluation and benchmarking

All experiments were managed with Hydra [22], enabling standardized configuration and reproducibility. Performance was evaluated using mean reciprocal rank (MRR), a standard information retrieval metric that measures how highly the correct item appears in a ranked list. Formally, for each drug *i*, let *r*_*i*_ denote the rank of the correct item. The reciprocal rank is defined as 1*/r*_*i*_, and the overall score is computed as the mean across all drugs:

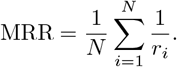

This formulation places the greatest weight on identifying the correct answer near the top of the ranking, while penalizing errors at lower positions only minimally. As such, MRR is well suited to our setting, where the value of SigSpace depends on surfacing the correct drug or MoA among the leading predictions derived from anonymized summaries.

Additionally, as an MoA prediction benchmark, we used a non-parametric k-nearest-neighbor (k-NN) classifier that assigns a drug’s MoA by comparing its transcriptional-response signature with those of other drugs measured under the same conditions. For each drug, we constructed binary gene-set features mirroring the preprocessing used before LLM summarization: for Vision, the top-*N* up-regulated and bottom-*N* down-regulated sets (*N* = 250); for GSEA, all gene sets passing a fraction gate (*p*_adj_ *<* 0.05 in a consistent direction in ≥ 25% of cell lines). Jaccard drug-drug distances were computed separately for the up- and down-regulated feature matrices and then averaged, excluding self-distances. Each drug’s MoA was predicted from its *k* nearest neighbors (*k* = 10, 15) by majority vote with closest-neighbor tie-break, and we report MRR from the induced MoA ranking.

## 3 Results

Table 1 summarizes the performance of different models and signature score combinations for the drug ranking and MoA ranking tasks. Higher MRR values indicate better ranking performance. The baseline is the expected MRR under random ranking (Supplementary Section 5).

**Table 1:** MRR of drug ranking and MoA ranking tasks.

| Model + input | Drug ranking MRR<br>(375 drugs) | MoA ranking MRR<br>(25 MoAs) |
| --- | --- | --- |
| DeepSeek-R1+ Vision | 0.021 | **0.218** |
| DeepSeek-R1 + GSEA | **0.026** | 0.198 |
| o4-mini + Vision | 0.021 | **0.214** |
| o4-mini + GSEA | **0.028** | 0.210 |
| Llama-4-Maverick-17B + Vision | 0.019 | 0.197 |
| Llama-4-Maverick-17B + GSEA | 0.017 | 0.187 |
| Baseline (expected MRR) | 0.017 | 0.153 |

Top 2 combinations per task are shown in bold

Most model-signature score combinations outperform the random ranking baseline. GSEA scores generally yield higher performance on the drug ranking task, whereas Vision scores perform better on the MoA ranking task. DeepSeek-R1 and o4-mini show stronger overall performance than Llama-4-Maverick-17B. These results underscore the dependence of performance on the LLM, the signature score input, and the specific prediction task.

In Figure 2, we show example MoA ranking results for o4-mini with Vision scores, compared against a random baseline. Most MoAs (16/25, orange bars) outperform the baseline, with no clear bias based on the number of drugs per MoA (ranging from 27 for DNA synthesis/repair inhibitors to 3 for RAF inhibitors and other smaller categories). Supplementary Figure 2 presents drug ranking results for o4-mini with GSEA scores, where true drugs rank better than random, with over 15% appearing in the top 50 predictions. These top-50 predictions were further enriched for drugs sharing the same MoA as the true drug (Supplementary Figure 3).

**Figure 2:**
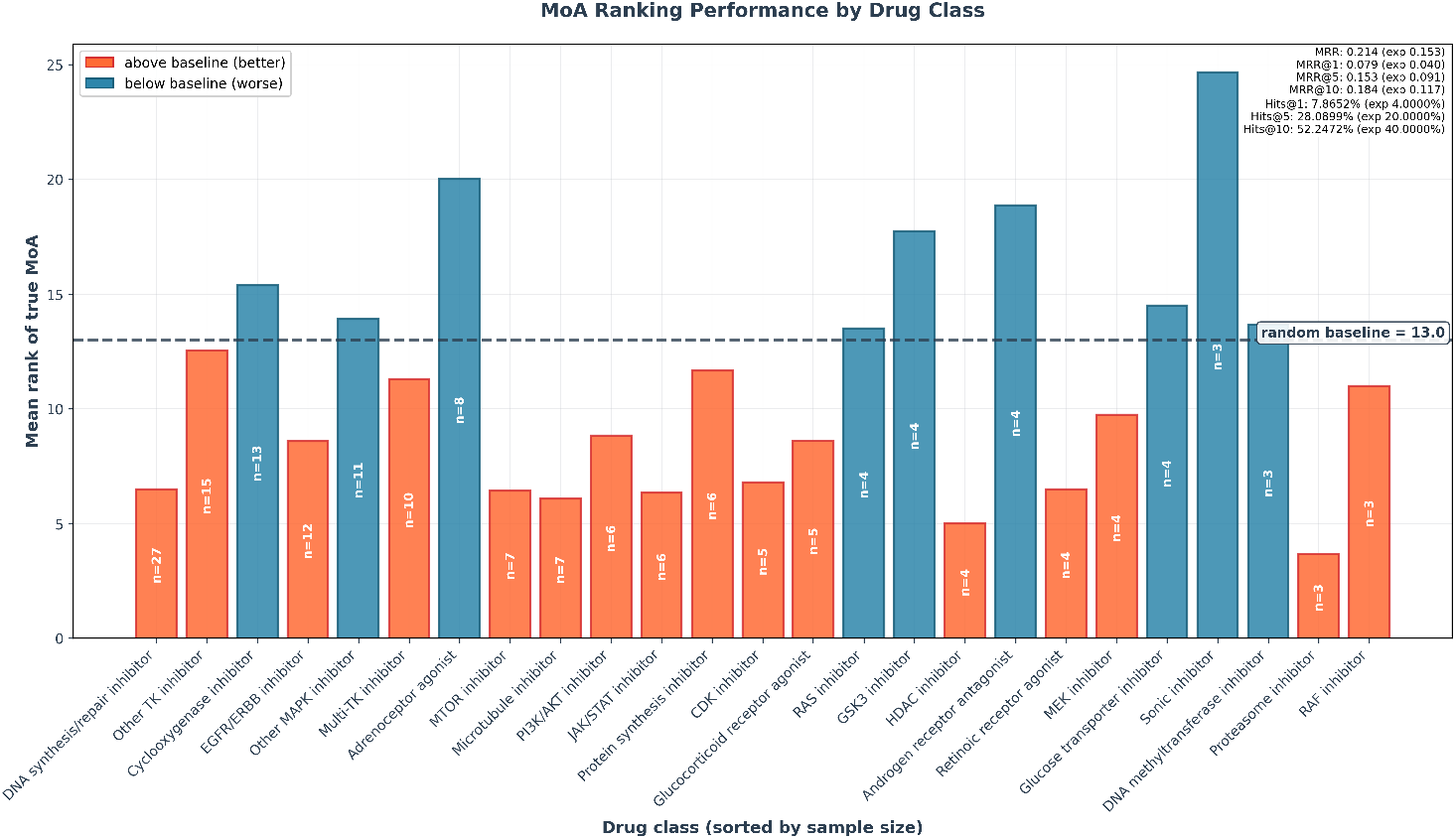
MoA ranking performance using o4-mini and Vision scores from Tahoe-100M. Each bar represents an MoA class, sorted in descending order by sample size (i.e., number of drugs with that MoA), with the height showing the mean rank of the true MoA. Orange bars indicate performance above the random baseline (expected mean rank under random ranking), while blue bars indicate performance below baseline. MRR@*k* rewards precision of ranking within the top-*k*, while Hits@*k* measures whether the correct answer appears in the top-*k* predictions.

To further contextualize MoA ranking performance, we implemented a k-NN classifier that assigns a drug’s MoA by comparing its transcriptional-response signature to those of other drugs. This yielded MRRs of 0.328 for *k* = 15 and 0.338 for *k* = 10 with Vision scores and 0.213 for *k* = 15 and 0.226 for *k* = 10 with GSEA scores. Unlike the LLM, which must infer MoAs from first principles, k-NN relies on direct similarity matching, a less demanding task since transcriptional responses naturally cluster drugs with related MoAs, even though external reference knowledge may be insufficient to infer them. In addition, the k-NN classifier performance is more biased by the number of drugs per MoA class, in comparison to the LLM predictions (Supplementary Figure 4). Lastly, the k-NN approach cannot be applied to the drug ranking task, as it requires prior knowledge of the drug’s MoA to perform majority voting. Nevertheless, this baseline serves as a useful positive control, providing intuition for the difficulty of MoA prediction and contextualizing the LLM-based performance.

Figure 3 exemplifies the summarization process, with excerpts for the drug Trametinib, an approved MEK1/2 inhibitor, capturing its known ability to upregulate the interferon-alpha response and antigen presentation [23] as well as downregulate RAS/MAPK oncogenic signaling and pathways related to the epithelial-mesenchymal transition [24]. Additional examples are provided for the microtubule-interfering agents Paclitaxel and Docetaxel, which induce upregulation of the mitotic spindle and related pathways, consistent with spindle-assembly checkpoint activation and M-phase cell cycle arrest (Supplementary Figure 5).

**Figure 3:**
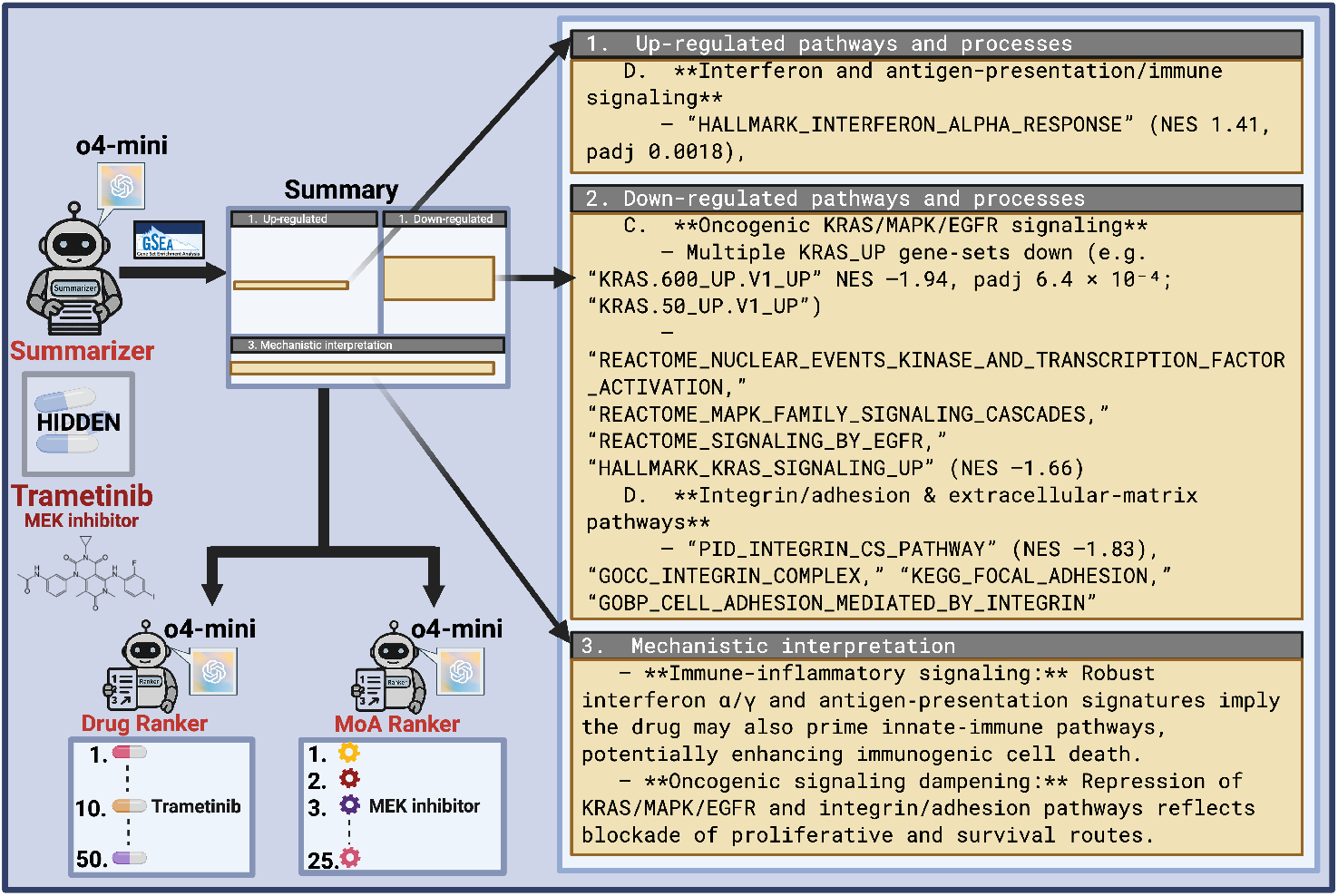
Summary and results for Trametinib using SigSpace. The system reasons over GSEA scores, uses o4-mini to generate a human-readable summary of the perturbed pathways, and then returns ranked predictions for either drug identity or MoA, where the drug identity is hidden during ranking. The predicted ranks were 10 for the drug and 3 for the MoA.

We additionally investigated 34 drugs from two very well-characterized MoAs, DNA synthesis/repair inhibitors and microtubule inhibitors, which are expected to produce particularly strong response signatures since they often correspond to commonly used cancer chemotherapeutics. We observed higher MRRs than when considering all drugs. For this group, we additionally evaluated if injecting information about cell line-specific responses produced any performance differences, but we did not observe any consistent trend (Table 2). This suggests that incorporating cell-line-specific metadata did not improve the LLM’s ability to rank drug MoAs in this setting.

**Table 2:** MoA ranking performance with and without cell-line metadata, benchmarked on a subset of drugs: DNA synthesis/repair inhibitors (27) and microtubule inhibitors (7)

| Model + input | MoA MRR without metadata<br>(25 MoAs, 34 drugs) | MoA MRR with metadata<br>(25 MoAs, 34 drugs) |
| --- | --- | --- |
| DeepSeek-R1 + Vision | **0.412** | **0.426** |
| DeepSeek-R1 + GSEA | 0.305 | 0.303 |
| o4-mini + Vision | 0.383 | 0.272 |
| o4-mini + GSEA | 0.279 | 0.262 |
| Llama-4-Maverick-17B + Vision | 0.298 | 0.289 |
| Llama-4-Maverick-17B + GSEA | 0.251 | 0.268 |
| Baseline (expected MRR) | 0.153 | 0.153 |

Finally, we investigated error patterns for both the drug ranking and MoA ranking tasks by visualizing true vs. predicted rank heatmaps (Supplementary Figures 6, 7). We noticed that MTOR inhibitor, proteasome inhibitor, and protein synthesis inhibitor were consistently and incorrectly predicted as top MoAs, and drugs in these MoA classes were similarly incorrectly predicted as top drugs (4EGI-1, Bortezomib, Everolimus, Ixazomib, Rapamycin, Sapanisertib, etc.). This does not appear to be an artifact of LLM summarization, but rather is driven by the frequent presence of response signatures related to protein synthesis and ribosomes, which are captured in a large fraction of the resulting summaries (Supplementary Table 1). This issue may be addressed in a future iteration by removing or flagging more promiscuous signatures.

## 4 Conclusions

This study demonstrates that LLMs offer valuable support for the summarization and interpretation of transcriptional response signatures to drug perturbations in cancer cell lines. Performance for predictive tasks such as drug identity and MoA reconstruction, while modest, was better than a random baseline and was encouraging given the complexity of these predictive tasks from transcriptional data alone. Performance varied systematically across LLM models and gene signature methods, and showed overall high results for well-characterized MoAs such as DNA synthesis/repair inhibitors and microtubule-interfering perturbations. Finally, the summaries are human-readable and provide literature-consistent insights into drug mechanisms, supporting hypothesis generation and interpretation of gene-signature results. Additional future work should focus on improving summarization performance, benchmarking prediction performance for different output summary sizes, and evaluating additional datasets and contextual information.

## 5 Supplementary material

**Supplementary Figure 1:**
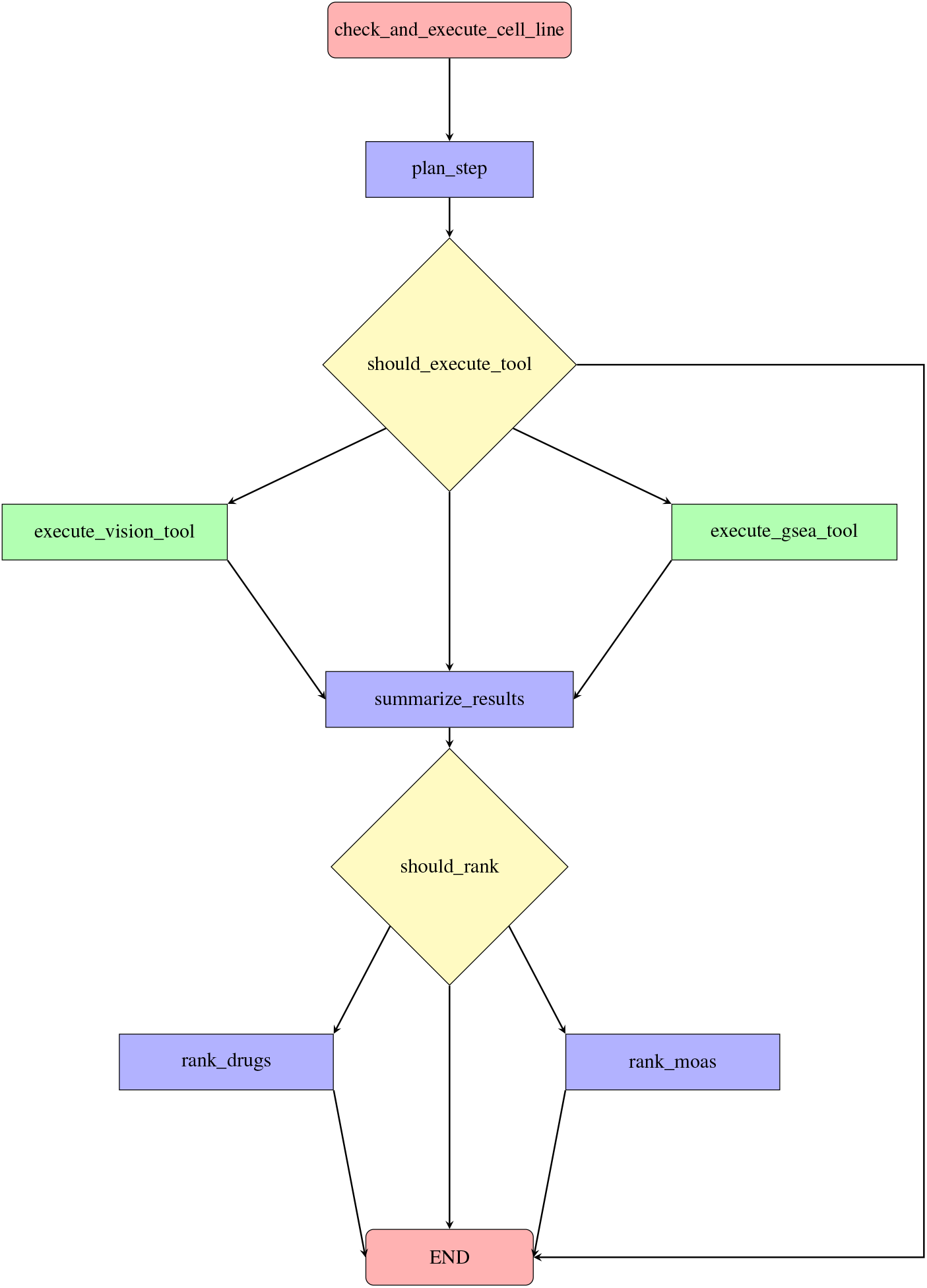
SigSpace’s computational workflow. The process begins with cell-line validation, followed by conditional branching for tool execution (Vision or GSEA score filtering), result summarization, and ranking of drugs or MoAs before termination.

### Baseline MRR under random ranking

Let *N* be the number of drugs (queries) and *M* the number of candidate items (i.e., drugs for the drug ranking task or MoAs for the MoA ranking task).

For drug *i*, let *r*_*i*_ denote the rank of the correct item, and define:

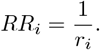

Then, the MRR is:

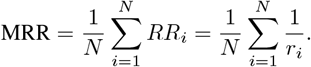

For random ranking, *r*_*i*_ is uniformly distributed over 1, 2, …, *M* with Pr(*r*_*i*_ = *r*) = 1/*M*. Thus, for each drug *i*:

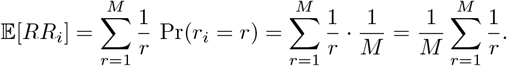

By linearity of expectation:

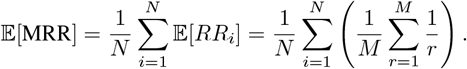

Since the inner term does not depend on *i*, this simplifies to:

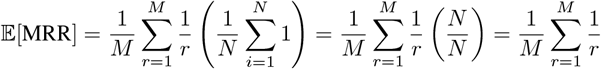

The summation is the *M*-th harmonic number:

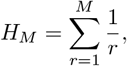

so the random baseline for MRR is:

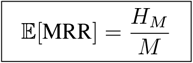

### Task-specific examples

- For MoA ranking, we have *M* = 25 MoA classes and *H*_25_ ≈ 3.816, hence:

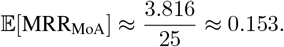
- For drug ranking, we have *M* = 375 drugs and *H*_375_ ≈ 6.505, hence:

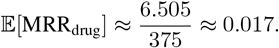

---

**Supplementary Figure 2:**
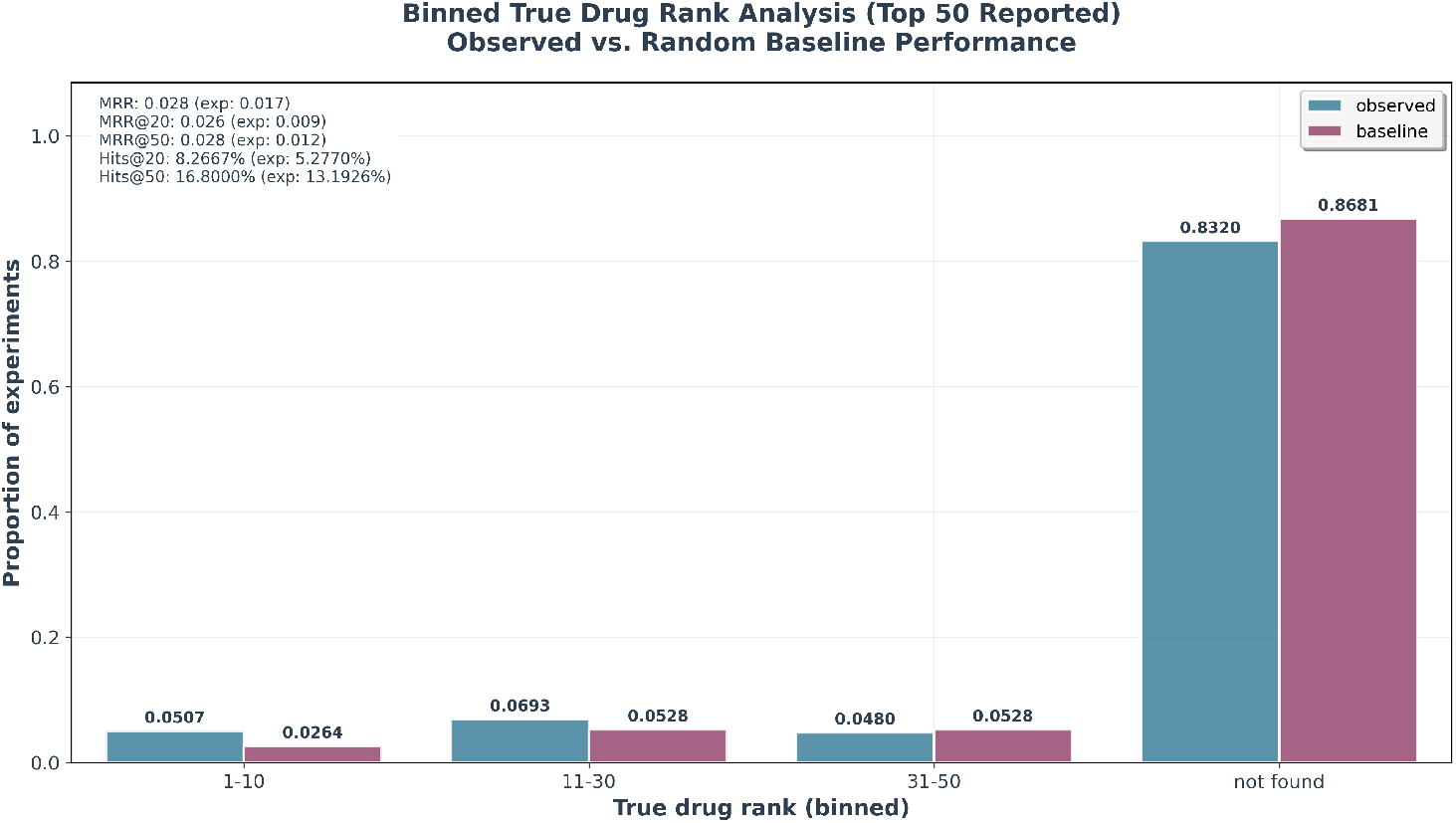
Drug ranking performance using o4-mini and GSEA scores from Tahoe-100M. The random baseline represents the expected performance if drugs were ranked uniformly at random.

**Supplementary Figure 3:**
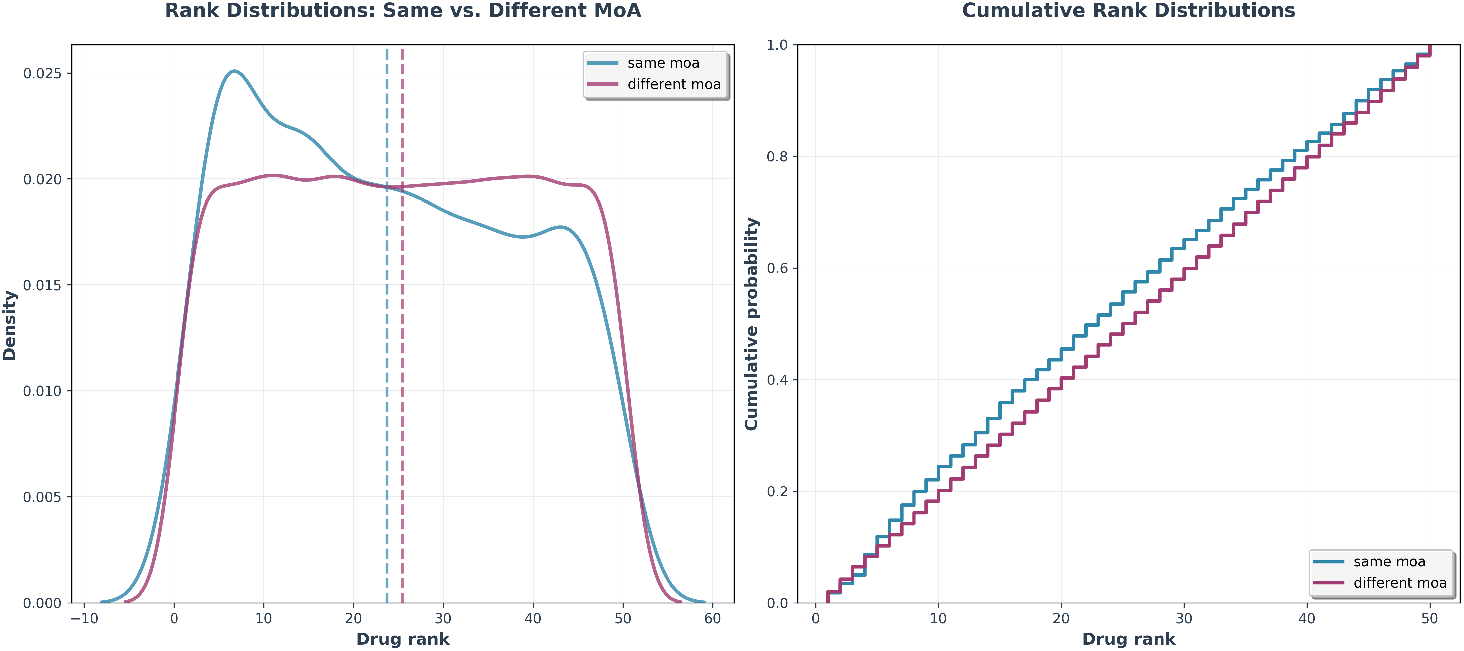
MoA analysis of drug ranking performance using o4-mini and GSEA scores from Tahoe-100M. Rank distributions are shown for drugs sharing the same MoA as the true drug vs. those with different MoAs, with density plots (left) and cumulative distributions (right): drugs with the same MoA as the true drug tend to have better ranks, illustrating the model’s ability to distinguish between related and unrelated drug classes.

**Supplementary Figure 4:**
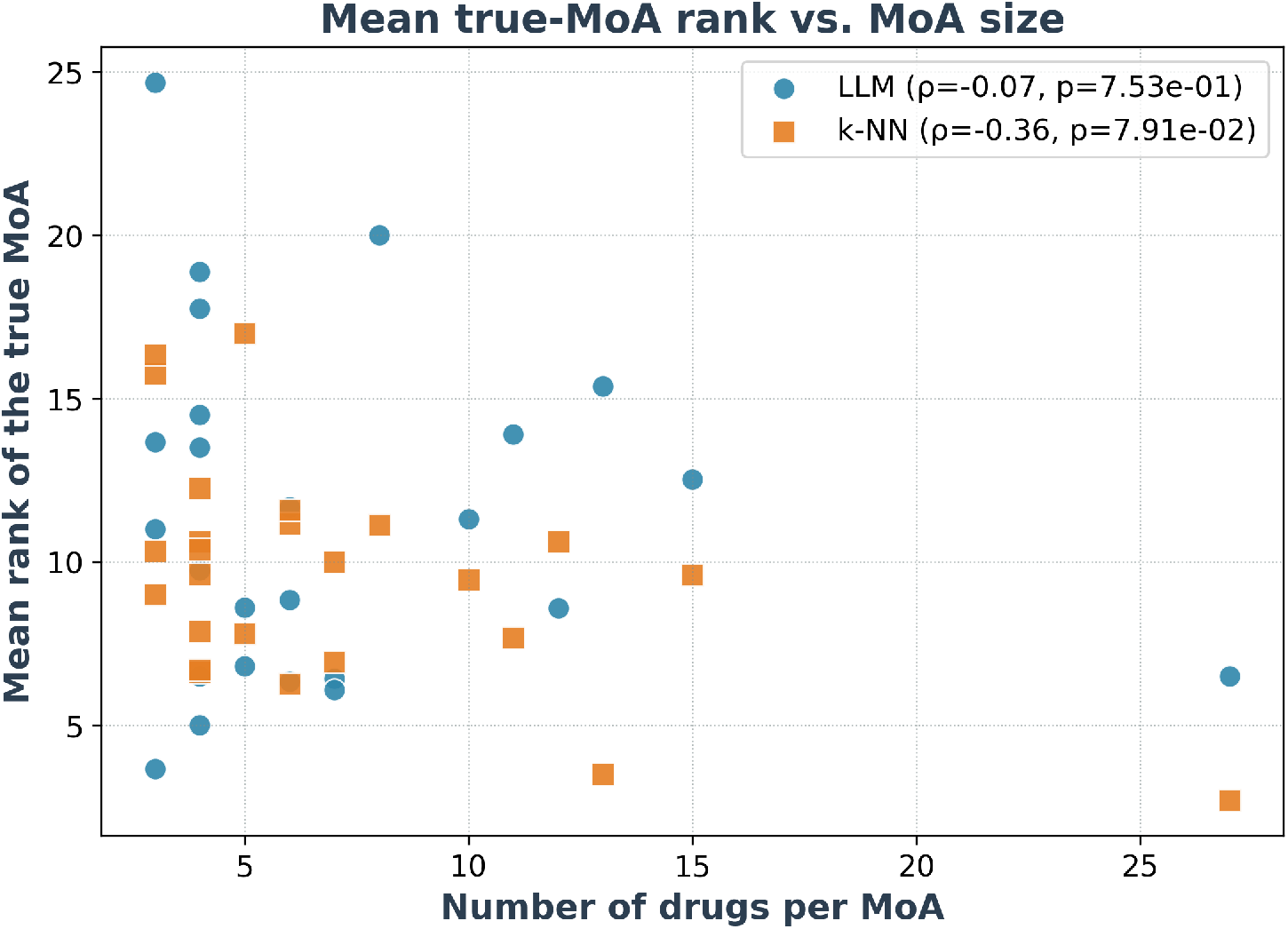
Scatterplot of the true MoA mean rank by the number of drugs per MoA, for the LLM (o4-mini) and k-NN with Vision scores as input, showing that for the k-NN there is a significant correlation between ranking performance and MoA’s drug numerosity.

**Supplementary Table 1:** Percentage of 375 drug response summaries containing “protein synthesis” or “ribosome” keywords (case-insensitive)

| Model + input | Query | % of drug ranking summaries | % of MoA ranking summaries |
| --- | --- | --- | --- |
| DeepSeek-R1 + GSEA | protein synthesis | 87.9 | 85.9 |
| DeepSeek-R1 + Vision | protein synthesis | 49.9 | 70.4 |
| o4-mini + GSEA | protein synthesis | 71.2 | 67.5 |
| o4-mini + Vision | protein synthesis | 47.5 | 50.1 |
| Llama-4-Maverick-17B + GSEA | protein synthesis | 83.5 | 84.8 |
| Llama-4-Maverick-17B + Vision | protein synthesis | 65.9 | 66.4 |
| DeepSeek-R1 + GSEA | ribosome | 85.0 | 87.2 |
| DeepSeek-R1 + Vision | ribosome | 50.9 | 71.2 |
| o4-mini + GSEA | ribosome | 90.7 | 90.4 |
| o4-mini + Vision | ribosome | 78.7 | 78.9 |
| Llama-4-Maverick-17B + GSEA | ribosome | 81.1 | 81.9 |
| Llama-4-Maverick-17B + Vision | ribosome | 63.5 | 64.0 |

**Supplementary Figure 5:**
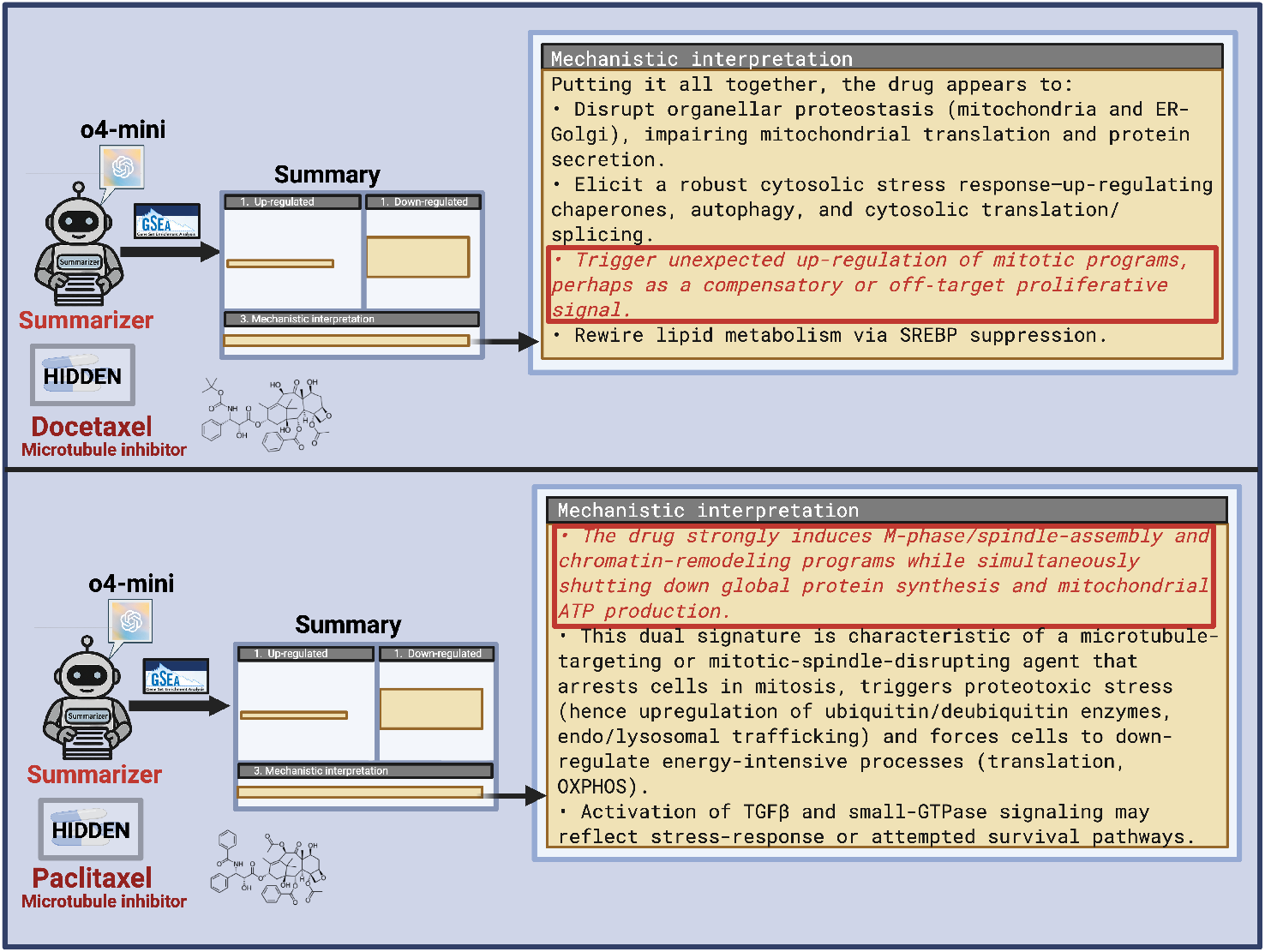
SigSpace-generated summaries for the microtubule inhibitors Docetaxel and Paclitaxel. Based on GSEA scores and o4-mini, the outputs highlight consistent perturbations in mitotic spindle-associated pathways, reflecting activation of spindle-assembly checkpoints and disruption of normal cell-cycle progression.

**Supplementary Figure 6:**
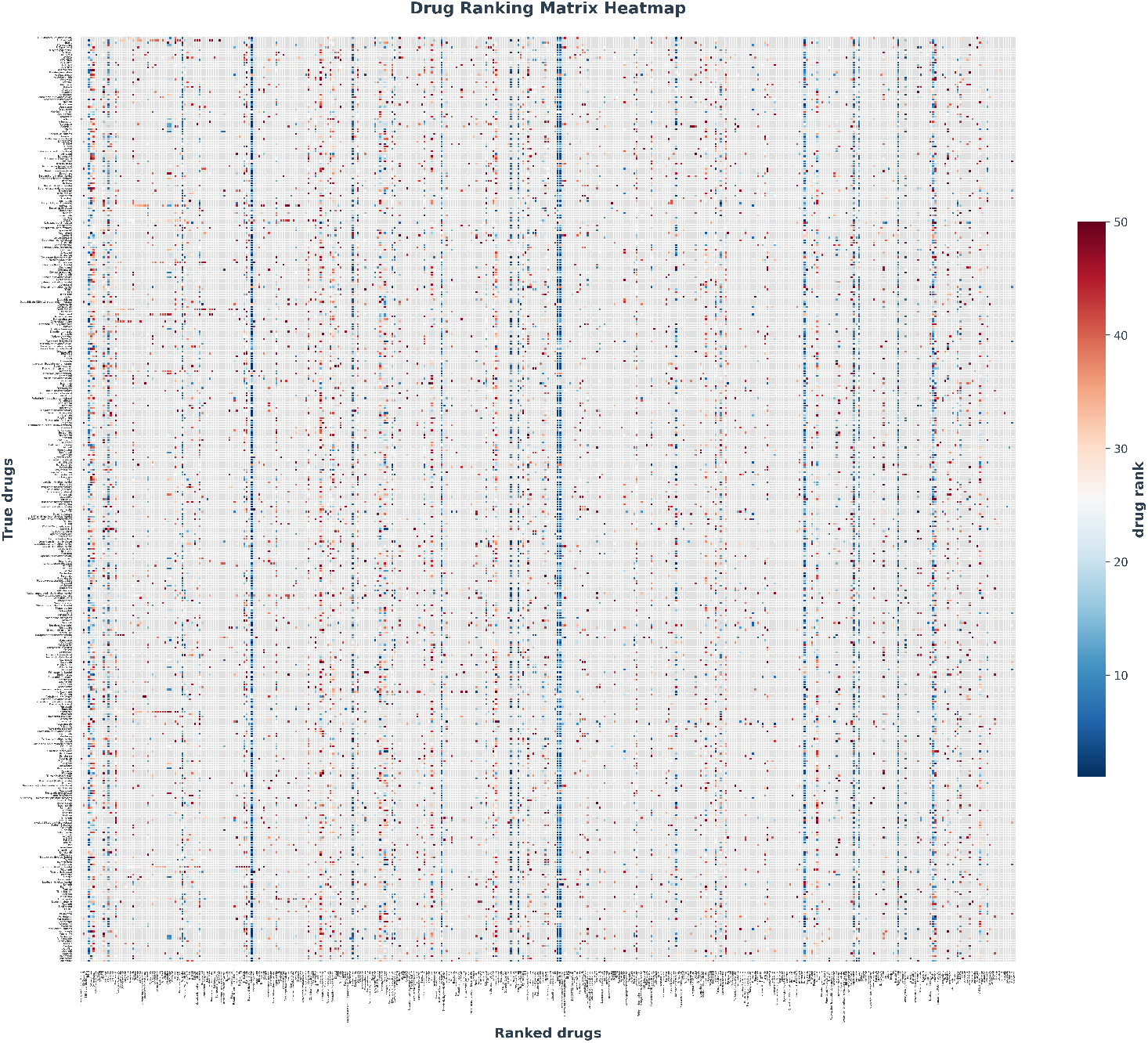
Drug ranking matrix heatmap using o4-mini and GSEA scores from Tahoe-100M. Rows represent true drugs and columns represent ranked drugs. Color intensity indicates rank position (dark blue = better rank), with gray cells showing drugs not within the top 50. Vertical blue striping corresponds to drugs that disproportionately achieve better ranks, regardless of true drug characteristics (examples include Bortezomib, Everolimus, Harringtonine, Ixazomib, etc.).

**Supplementary Figure 7:**
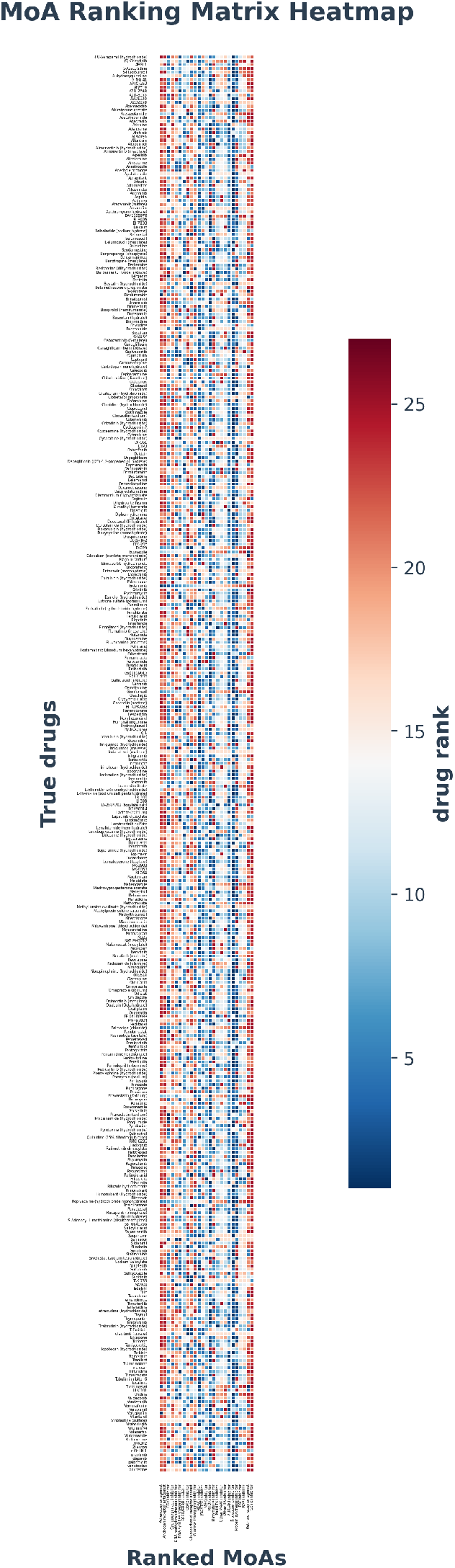
MoA ranking matrix heatmap using o4-mini and Vision scores from Tahoe-100M. Rows represent true drugs and columns represent ranked MoAs. Color intensity indicates rank position (dark blue = better rank). Vertical blue striping corresponds to MoAs that disproportionately achieve better ranks, regardless of true drug characteristics (examples include MTOR inhibitor, proteasome inhibitor, protein synthesis inhibitor, etc.).

## Footnotes

2 https://github.com/tahoebio/sigspace.git

## References

[1] Shanghua Gao, Ada Fang, Yepeng Huang, Valentina Giunchiglia, Ayush Noori, Jonathan Richard Schwarz, Yasha Ektefaie, Jovana Kondic, and Marinka Zitnik. Empowering biomedical discovery with ai agents. Cell, 187(22):6125–6151, 2024.

[2] Kexin Huang, Serena Zhang, Hanchen Wang, Yuanhao Qu, Yingzhou Lu, Yusuf Roohani, Ryan Li, Lin Qiu, Gavin Li, Junze Zhang, et al. Biomni: A general-purpose biomedical ai agent. biorxiv, pages 2025–05, 2025.

[3] Jinyuan Fang, Yanwen Peng, Xi Zhang, Yingxu Wang, Xinhao Yi, Guibin Zhang, Yi Xu, Bin Wu, Siwei Liu, Zihao Li, Zhaochun Ren, Nikos Aletras, Xi Wang, Han Zhou, and Zaiqiao Meng. A comprehensive survey of self-evolving ai agents: A new paradigm bridging foundation models and lifelong agentic systems, 2025.

[4] Andres M Bran, Sam Cox, Oliver Schilter, Carlo Baldassari, Andrew D White, and Philippe Schwaller. Chemcrow: Augmenting large-language models with chemistry tools, 2023.

[5] Justin Lamb, Emily D Crawford, David Peck, Joshua W Modell, Irene C Blat, Matthew J Wrobel, Jim Lerner, Jean-Philippe Brunet, Aravind Subramanian, Kenneth N Ross, et al. The connectivity map: using gene-expression signatures to connect small molecules, genes, and disease. Science, 313(5795):1929–1935, 2006.

[6] Sudeep Pushpakom, Francesco Iorio, Patrick A Eyers, K Jane Escott, Shirley Hopper, Andrew Wells, Andrew Doig, Tim Guilliams, Joanna Latimer, Christine McNamee, et al. Drug repurposing: progress, challenges and recommendations. Nature reviews Drug discovery, 18(1):41–58, 2019.

[7] Purvesh Khatri, Marina Sirota, and Atul J Butte. Ten years of pathway analysis: current approaches and outstanding challenges. PLoS computational biology, 8(2):e1002375, 2012.

[8] Jüri Reimand, Ruth Isserlin, Veronique Voisin, Mike Kucera, Christian Tannus-Lopes, Asha Rostamianfar, Lina Wadi, Mona Meyer, Jeff Wong, Changjiang Xu, et al. Pathway enrichment analysis and visualization of omics data using g: Profiler, gsea, cytoscape and enrichmentmap. Nature protocols, 14(2):482–517, 2019.

[9] Alexander Novikov, Ngân Vũ, Marvin Eisenberger, Emilien Dupont, Po-Sen Huang, Adam Zsolt Wagner, Sergey Shirobokov, Borislav Kozlovskii, Francisco J. R. Ruiz, Abbas Mehrabian, M. Pawan Kumar, Abigail See, Swarat Chaudhuri, George Holland, Alex Davies, Sebastian Nowozin, Pushmeet Kohli, and Matej Balog. Alphaevolve: A coding agent for scientific and algorithmic discovery, 2025.

[10] Kyle Swanson, Wesley Wu, Nash L. Bulaong, John E. Pak, and James Zou. The virtual lab of ai agents designs new sars-cov-2 nanobodies. Nature, July 2025.

[11] Ali Essam Ghareeb, Benjamin Chang, Ludovico Mitchener, Angela Yiu, Caralyn J. Szostkiewicz, Jon M. Laurent, Muhammed T. Razzak, Andrew D. White, Michaela M. Hinks, and Samuel G. Rodriques. Robin: A multi-agent system for automating scientific discovery, 2025.

[12] Sizhe Liu, Yizhou Lu, Siyu Chen, Xiyang Hu, Jieyu Zhao, Yingzhou Lu, and Yue Zhao. Drugagent: Automating ai-aided drug discovery programming through llm multi-agent collaboration, 2025.

[13] Namkyeong Lee, Edward De Brouwer, Ehsan Hajiramezanali, Tommaso Biancalani, Chanyoung Park, and Gabriele Scalia. Rag-enhanced collaborative llm agents for drug discovery, 2025.

[14] Marcin P Joachimiak, J Harry Caufield, Nomi L Harris, Hyeongsik Kim, and Christopher J Mungall. Gene set summarization using large language models. ArXiv, pages arXiv–2305, 2024.

[15] Zhizheng Wang, Qiao Jin, Chih-Hsuan Wei, Shubo Tian, Po-Ting Lai, Qingqing Zhu, Chi-Ping Day, Christina Ross, Robert Leaman, and Zhiyong Lu. Geneagent: self-verification language agent for gene-set analysis using domain databases. Nature Methods, 22(8):1677–1685, July 2025.

[16] Kaspar Märtens, Marc Boubnovski Martell, Cesar A Prada-Medina, and Rory Donovan-Maiye. Langpert: Llm-driven contextual synthesis for unseen perturbation prediction. In ICLR 2025 Workshop on Machine Learning for Genomics Explorations, 2025.

[17] Jesse Zhang, Airol A Ubas, Richard de Borja, Valentine Svensson, Nicole Thomas, Neha Thakar, Ian Lai, Aidan Winters, Umair Khan, Matthew G Jones, et al. Tahoe-100m: A gigascale single-cell perturbation atlas for context-dependent gene function and cellular modeling. BioRxiv, pages 2025–02, 2025.

[18] Aravind Subramanian, Pablo Tamayo, Vamsi K Mootha, Sayan Mukherjee, Benjamin L Ebert, Michael A Gillette, Amanda Paulovich, Scott L Pomeroy, Todd R Golub, Eric S Lander, et al. Gene set enrichment analysis: a knowledge-based approach for interpreting genome-wide expression profiles. Proceedings of the National Academy of Sciences, 102(43):15545–15550, 2005.

[19] David DeTomaso, Matthew G Jones, Meena Subramaniam, Tal Ashuach, Chun J Ye, and Nir Yosef. Functional interpretation of single cell similarity maps. Nature communications, 10(1):4376, 2019.

[20] Alexey Sergushichev. fgsea, 2017.

[21] Harrison Chase. Langchain: Build context-aware reasoning applications, 2022. Open source framework for building LLM-powered applications.

[22] Omry Yadan. Hydra - a framework for elegantly configuring complex applications. Github, 2019.

[23] Lauren Dennison, Aditya A Mohan, and Mark Yarchoan. Tumor and systemic immunomodulatory effects of mek inhibition. Current oncology reports, 23(2):23, 2021.

[24] Emily N Arner, Wenting Du, and Rolf A Brekken. Behind the wheel of epithelial plasticity in kras-driven cancers. Frontiers in oncology, 9:1049, 2019.

